# Introgression, hominin dispersal and megafaunal survival in Late Pleistocene Island Southeast Asia

**DOI:** 10.1101/2020.07.24.219048

**Authors:** João C. Teixeira, Guy S. Jacobs, Chris Stringer, Jonathan Tuke, Georgi Hudjashov, Gludhug A. Purnomo, Herawati Sudoyo, Murray P. Cox, Ray Tobler, Chris S.M. Turney, Alan Cooper, Kristofer M. Helgen

## Abstract

The hominin fossil record of Island Southeast Asia (ISEA) indicates that at least two endemic ‘super-archaic’ species – *Homo luzonensis* and *H. floresiensis* – were present around the time anatomically modern humans (AMH) arrived in the region >50,000 years ago. Contemporary human populations carry signals consistent with interbreeding events with Denisovans in ISEA – a species that is thought to be more closely related to AMH than the super-archaic endemic ISEA hominins. To query this disparity between fossil and genetic evidence, we performed a comprehensive search for super-archaic introgression in >400 modern human genomes. Our results corroborate widespread Denisovan ancestry in ISEA populations but fail to detect any super-archaic admixture signals. By highlighting local megafaunal survival east of the Wallace Line as a potential signature of deep, pre-*H. sapiens* hominin-faunal interaction, we propose that this understudied region may hold the key to unlocking significant chapters in Denisovan prehistory.

## Main Text

Island Southeast Asia (ISEA) hosts a unique fossil record of hominin presence throughout the Pleistocene^1^. *Homo erectus* has a deep history in the region, from the early Pleistocene until ~108ka^2^, and at least two additional endemic species are known to have survived in ISEA until, or close to, the arrival of anatomically modern humans (AMH) >50 thousand years ago (ka)^3–6^: *H. floresiensis* on Flores, in the Lesser Sundas^7,8^, and *H. luzonensis* on Luzon, in the northern Philippines^9^. The phylogenetic placement of these two species remains an area of debate. Recent interpretations suggest that *H. floresiensis* is either a close relative of *H. erectus*, or instead represents a separate dispersal event out of Africa of an earlier diverging species of *Homo^7,10,11^*. The current classification of *H. luzonensis* is also uncertain; specimens demonstrate similarities in certain morphological traits with various hominin taxa including *Australopithecus*, Asian *H. erectus, H. floresiensis* and *H. sapiens^9^*.

Genetic evidence preserved in modern human genomes suggests that at least one additional hominin group likely inhabited ISEA at the time of AMH arrival. Present-day populations living in ISEA, New Guinea and Australia harbour significant genetic ancestry from Denisovans, a sister lineage to Neanderthals, for which the fossil record remains scarce and mostly limited to the eponymous cave in the Altai Mountains in Siberia^12,13^, along with a >160,000-year-old mandible found in the Tibetan Plateau^14^. Despite this geographically circumscribed record, the spatial and genetic patterns of Denisovan admixture in modern human populations point to several independent events of AMH-Denisovan interbreeding occurring during the migratory movements that brought AMH through ISEA into Sahul (the former continental landmass that connected New Guinea with Australia up to 8ka)^15–17^. These events may have involved different Denisovan populations living in ISEA, including the Philippines^18^, New Guinea^16^ and, potentially, Flores^17,19^.

The disparity between the lack of fossil evidence for Denisovan presence in ISEA, and the likelihood of AMH-Denisovan mixing in this region, poses an important outstanding question in hominin prehistory. A solution to this riddle would be found if *H. luzonensis* and/or *H. floresiensis* could be identified as the potential sources of the “Denisovan” contributions to modern human genomes in the region; however, this solution is not supported by current morphological interpretations. The anatomical attributes of both of these extinct ISEA hominin species are not closely identifiable with the few confirmed specimens of Denisovans from Altai and Tibet, leading paleoanthropologists to place them outside the clade comprising Denisovans, Neanderthals, and modern humans^7–11,20–22^. Moreover, morphological and archaeological data suggest that the lineages of H. *floresiensis* and H. *luzonensis* have very deep roots in the region, deeper than the estimated timescale for the emergence of the Denisovans^7–11,20–22^. Thus, the source of Denisovan introgression into modern human genomes in ISEA currently lacks a corresponding fossil record.

If *H. floresiensis* and *H. luzonensis* do represent super-archaic human relatives, it is possible that they also admixed with AMH populations in ISEA and subsequently this highly divergent genetic ancestry might survive in present-day ISEA populations. Signals of super-archaic admixture have been observed in Altai Denisovans^23^ and, potentially, in Andaman populations^24–26^, suggesting that additional super-archaic introgression may remain undetected in modern human genomes.

## Results

To address these questions and provide further insights into the hominin prehistory of ISEA, we implemented the most comprehensive search for introgressed super-archaic regions in modern human genomes performed to date. We searched a total of 426 genomes from across the world, including 214 individuals from Papuan and ISEA populations^16^ (Supplementary Table S1), for genomic signatures compatible with introgression from archaic hominins, which could reveal previously unknown introgressed DNA from *H. floresiensis*, *H. luzonensis* or other hypothetical late-surviving super-archaic hominin species. To detect blocks of introgressed super-archaic DNA, we extended the analytical pipeline reported by Jacobs et al.^16^ by including a recently published HMM detection method – which we call HMM_Archaic_ – along with the two methods used by Jacobs and colleagues; i.e. ChromoPainter (CP)^27^ and a Hidden Markov Model (HMM)^28,29^. Importantly, HMM_Archaic_ differs from CP and HMM in that it does not require a reference genome to guide the detection of introgressed DNA, making it suitable for identifying DNA from super-archaic groups for which no genome information currently exists. Accordingly, we were able to distinguish putative introgressed super-archaic blocks by running the three detection methods on all 426 genomes and only retaining those that did not overlap any of the Neanderthal and Denisovan blocks predicted by CP and/or HMM. We term the resulting set putative super-archaic sequences as residual_Archaic_ blocks (see Methods).

### No evidence for super-archaic introgression in AMH

Filtering the HMM_Archaic_ introgressed blocks overlapping Neanderthal- and Denisovan-introgressed tracts identified ~12.5Mb of residual_Archaic_ sequence per individual (Figure 1A). The amount of detected residual_Archaic_ sequence was consistent across worldwide populations, with a slightly higher amount found in East ISEA (~15Mb), and Papuan and Australian populations (~18Mb). In accordance with previous results, ISEA, Papuan, and Australian populations also had the largest amounts of Denisovan ancestry (reaching ~60Mb in Papuan and Australian genomes), meaning that these populations actually had the lowest proportion of residual_Archaic_ sequence relative to the total archaic ancestry observed across all analysed populations (Supplementary Figure S1). Our results indicate that super-archaic ancestry could potentially comprise a small but consistent amount of the genomic ancestry of modern human populations outside of Africa; however the current lack of support for demographic scenarios involving widespread super-archaic admixture in previous studies suggests that this global residual_Archaic_ signal is more likely a methodological artefact or a signal, ancient structure in human populations predating the out-of-Africa migration, or segregation of highly divergent AMH-derived sequences that were not detected in our African reference samples that result from incomplete lineage sorting or balancing selection^30^.

**Figure 1.**
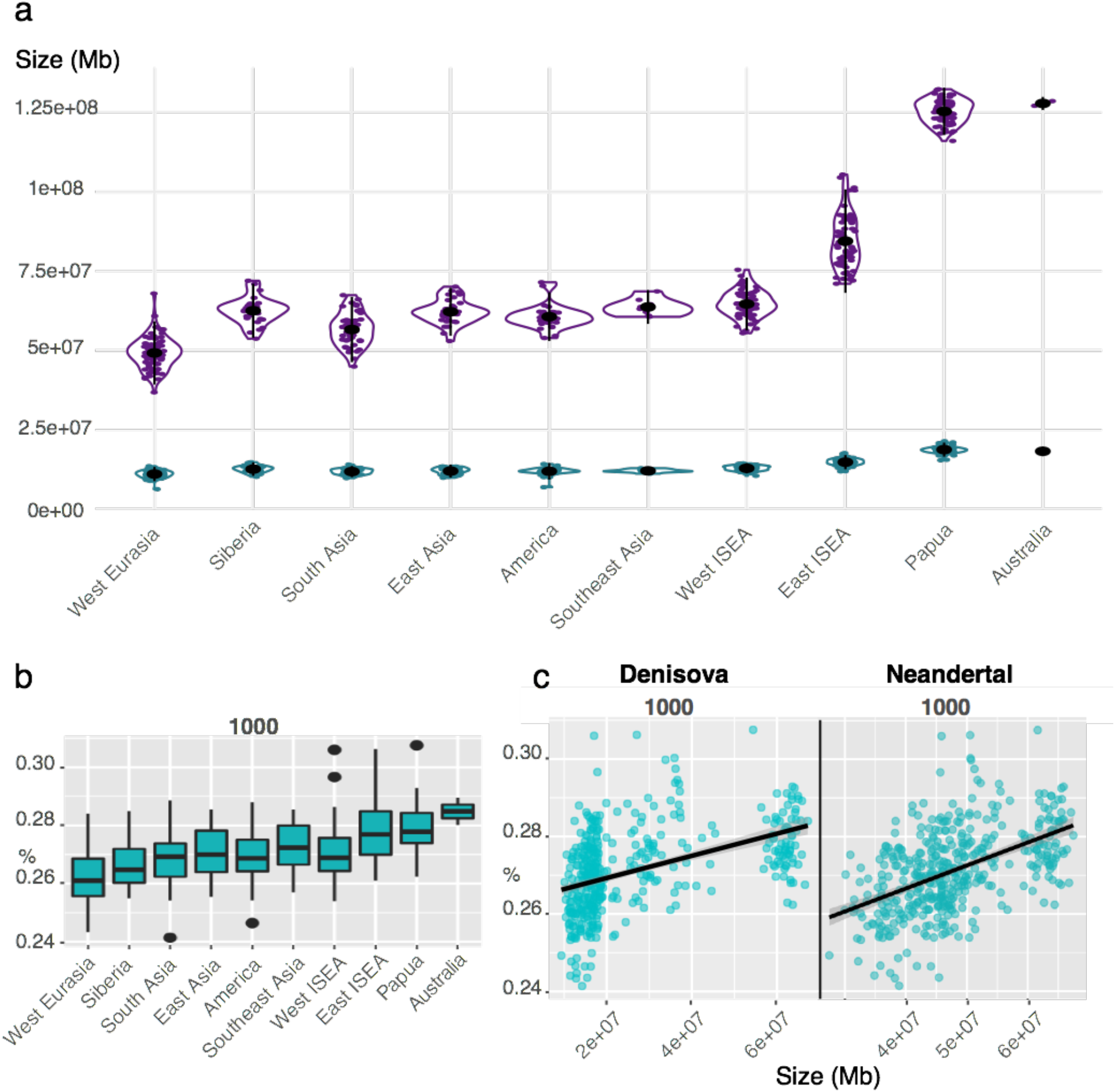
Introgression signals in extant populations across Island Southeast Asia. (a) Violin plots showing the cumulative amount (Mb) of Neanderthal and Denisovan ancestry (purple) estimated using HMM and residual_Archaic_ sequence (green) across different populations. Each dot represents a single sampled individual for a particular population. Within each violin plot, the population’s mean and 95% values of the distribution are shown as a black dot and vertical line, respectively. (b) The proportion of variants within residual_Archaic_ fragments that show mutation motifs compatible with super-archaic introgression [1000] per population. Each number on the string [1000] corresponds to the allelic states observed in [X, Denisovan, Neanderthal, Africa], where X is an individual from the test population (e.g. Australia), and 1 and 0 define derived and ancestral allelic states, respectively. (c) Scatter plot showing the association between the proportion of [1000] motifs within residual_Archaic_ fragments and the total amount of Denisovan (left) and Neanderthal (right) ancestry per individual.

Similarly, the additional ~2.5 to ~5Mb of residual_Archaic_ sequence observed in Papuan and Australian populations may represent a small but meaningful amount of super-archaic ancestry specific to this region, or instead simply reflect inter-population variation in the power of the statistical methods to detect Denisovan fragments or some other methodological artefact.

To further discriminate if the residual_Archaic_ blocks were truly introgressed super-archaic DNA, we searched for concordant signatures by investigating genetically distinct mutation motifs (i.e. allelic states) that are characteristic of introgressed super-archaic DNA within residual_Archaic_ blocks. Specifically, for each nucleotide position in a residual_Archaic_ block, we characterized the allelic state for the test individual (X), Denisova (D), Neanderthal (N), and an African individual (H) (see Methods). This resulted in a set of mutation motifs of the form [X, D, N, H], with patterns of the type [1000] and [0111] potentially indicative of super-archaic introgression signals. After enumerating these mutation motifs for all residual_Archaic_ blocks in each individual, we used generalised linear models to test if the proportion of motifs differed across the different worldwide populations, and computed *p*-values by contrasting the full model to a null model consisting of the intercept alone (see Methods).

The mutation motifs differed significantly between populations both when considering a linear model (ANOVA *p*-value 5.79×10^−224^) but not a multinomial logistic regression (where motifs are not independent as is assumed for the linear model; Figure 1B and Supplementary Figure S2). However, these differences are extremely subtle and correlate strongly with known archaic ancestry, suggesting a confounding effect (Figure 1C and Supplementary Figures S3-S6). For example, Papuan genomes show a slightly higher proportion of [1000] motifs (<2%) compared to other populations (Figure 1B and Supplementary Figure S2), but inter-individual variation is also high and we do not observe a similar increase in the proportion of the [0111] motif in the population, which is also expected under a scenario of super-archaic introgression (Supplementary Figure S2 and Methods).

While precise accounting for all motif count differences is non-trivial, likely explanations include the misclassification of alleles as either ancestral or derived, complex demographic histories, and the persistence of Neanderthal and Denisovan archaic signals amongst the residual_Archaic_ blocks that were not removed during the filtering step. For instance, the 2.5-5Mb extra residual_Archaic_ sequence observed in Papuans and Australians might have resulted from these populations having substantially more introgression from a Denisovan-like source that is highly divergent from the Altai Denisovan genome^16^. This may result in some of the more diverged blocks being detected by the reference-free HMM_Archaic_ scan, but not in the two methods that rely on reference genomes (i.e. CP and HMM). Indeed, while Denisovan and Neanderthal ancestry is positively correlated with the proportion of the [1000] motif across all populations, it is negatively correlated with the proportion of the [0111] motif (Supplementary Figures S3 and S4, respectively), which strongly suggests that differences in the proportion of these motifs is caused by unassigned Neanderthal and Denisovan ancestry within residual_Archaic_ blocks.

### Coalescent simulations support empirical observations

To rule out the possibility that the lack of evidence for super-archaic introgression into modern humans was due to a lack of power in our experimental design, we used the coalescent software *msprime^31^* to simulate Aboriginal Australian and Papuan histories under an empirically-informed demographic model^32^. These simulations included separate Neanderthal and Denisovan admixture events along with differing amounts of super-archaic introgression (2%, 1%, 0.1% and 0%) in the common ancestral population of Australo-Papuans (see Methods). We then applied our full analytical pipeline to these simulated genomic datasets to detect super-archaic blocks and quantified the power and false discovery rate for the different levels of super-archaic introgression.

Our simulation results demonstrate that HMM_Archaic_ can confidently detect super-archaic blocks even in scenarios with extremely low levels of super-archaic ancestry – with true positive rates (TPR) ranging from ~50% to ~95% for models with 0.1% and 2% super-archaic ancestry, respectively (Figure S9) – while maintaining extremely low false positive rates (Figure S10).

The amount of residual_Archaic_ sequences detected per individual in the 0.1% and 0% super-archaic introgression models (~20Mb – Figure 2a) is strikingly close to that observed in the Papuan and Australian empirical data (~18Mb – Figure 1A). For these models, the majority of the residual_Archaic_ signal originates from Neanderthal and Denisovan introgression that went undetected by CP and HMM (Figure 2 and Supplementary Figure S12). In contrast, the 1% and 2% super-archaic introgression models detect at least 2 times more residual_Archaic_ sequence per individual than empirical estimates (~33Mb and ~47Mb, respectively – Figure 2), which was primarily caused by an inflation in the number of super-archaic blocks. Interestingly, the detection of Neanderthal and Denisovan blocks by HMM_Archaic_ is severely affected with increasing amounts of super-archaic ancestry, as the power of this method is proportionate to the divergence between the introgressing archaic population and the outgroup human population (see Methods).

**Figure 2.**
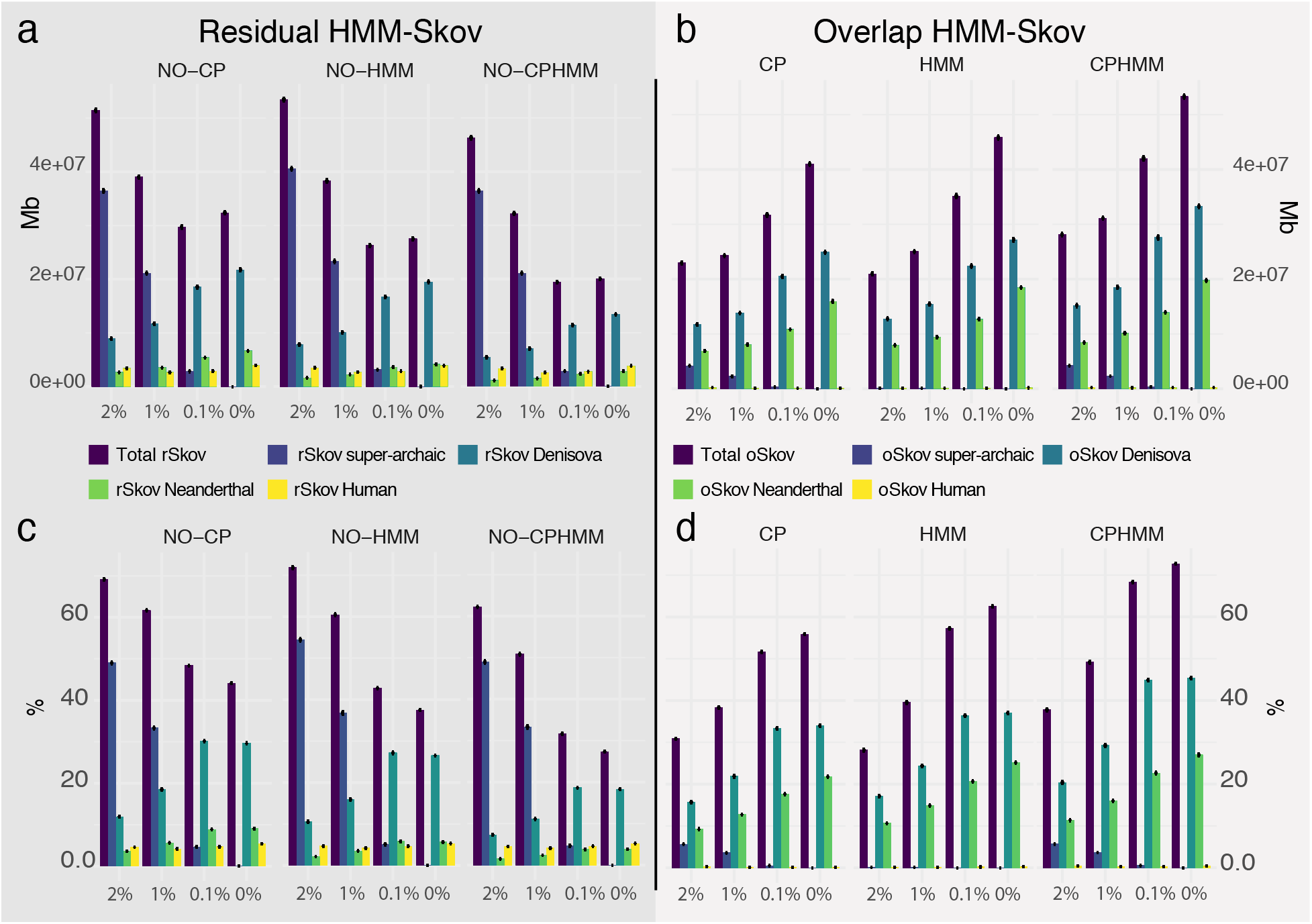
Results from coalescent simulations exploring the detection of archaic hominin introgressed sequences. a) residual_Archaic_ after removing Neanderthal and Denisovan regions detected by *CP*, *HMM* and *CP+HMM*. The total residual_Archaic_ and the proportion of residual_Archaic_ that overlap simulated archaic regions for different species is shown from left to right, together with the amount of residual_Archaic_ that overlaps ‘Human’ (i.e. non-archaic) regions. Different simulation models of super-archaic introgression are shown in the x-axis from left to right. b) Overlap between regions inferred as ‘archaic’, showing the concordance between HMM_Archaic_ and the other two methods (overlapArchaic). c) The proportion of residual_Archaic_ sequence over the total amount of HMM_Archaic_ inferred to be ‘archaic’. d) The proportion of overlapArchaic out of the total amount of HMM_Archaic_.

Similarly, the mutational motifs observed in 0.1% and 0% super-archaic introgression models provide a closer fit to the empirical data than higher levels of super-archaic introgression. For instance, the [1000] and [0111] mutational motifs comprise ~27% and ~6% on average in the empirical data, compared to ~26% and ~6.5% for the 0.1% model, and ~22.5% and ~4% for the 0% model (Figure S13). The close fit of the 0% and 0.1% models to our empirical observations provide strong support for there being little to no introgressed super-archaic sequences in non-African human genomes (Figure 2 and Supplementary Figure S12).

### Location of ISEA Denisovan groups and hominin interactions

Despite the late survival of multiple hominin species in ISEA, our results demonstrate that detectable super-archaic ancestry is absent from genomes of present-day non-African human populations. While genetic data can provide a reasonable proxy for inferring historical locations of these AMH-Denisovan encounters^17^, making more robust conclusions is complicated by the absence of any fossil specimens in ISEA currently attributable to Denisovan lineages. Hence, to further explore the likely locations of the AMH-Denisovan contact points in ISEA, we considered the possibility that the current distribution of endemic megafauna on islands in the region may offer clues to the past occurrence of hominins prior to the arrival of *H. sapiens*.

During the Pleistocene, diverse assemblages of very large vertebrates (megafauna) characterized the terrestrial biotas of most areas of the world, including the Sunda Shelf (the continental extension of mainland Asia, including the large islands of Sumatra, Java, and Borneo), the Sahul Shelf (Australia, Tasmania, and New Guinea), and the islands in between that were never connected by land to either continental shelf during the Pleistocene (Wallacea and the Philippines; Figure 3). Terrestrial animals larger than modern humans that occurred on the Sunda Shelf during the late Pleistocene, when *H. erectus* occupied the region, include the elephant *Elephas maximus*, the rhinos *Dicerorhinus sumatrensis* and *Rhinoceros sondaicus*, the tapir *Acrocodia indica*, the wild cattle *Bos javanicus*, the deer *Rusa* spp. and *Axis* spp., pigs *Sus*, orangutans *Pongo* spp., and the tiger *Panthera tigris*, as well as large pythons, *Python reticulata*.

**Figure 3.**
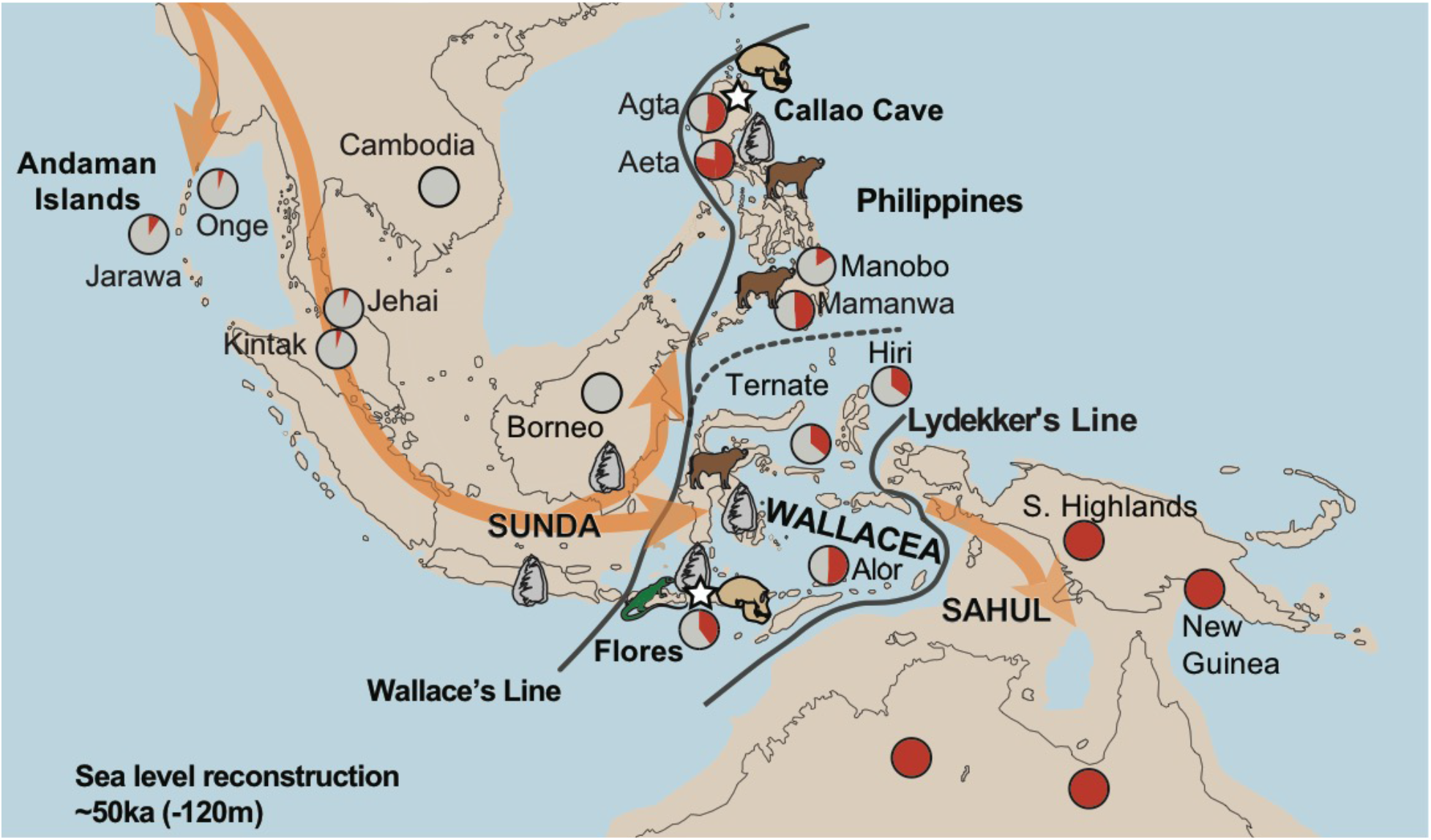
Hominin occupation and megafauna survival in Island Southeast Asia at the time of modern human arrival. Confirmed presence of *H. floresiensis* and *H. luzonensis* depicted by skull icons; regions with known artefacts associated with hominin presence are depicted by the stone tool icons; extant native megafauna east of the Wallace Line is depicted by the buffalo icon (representing mammals—*Bubalus, Rusa, Sus*, and *Babirusa*) in the northern and southern Philippines and Sulawesi, and Komodo dragon icon on Flores and satellites. Inferred hominin presence covers the entry routes into Sahul, indicated by the orange arrows. The estimated Denisovan ancestry in modern populations is shown in red in the pie charts, relative to that observed in Australo-Papuan genomes. All populations containing large amounts of Denisovan ancestry are found east of Wallace’s Line. Major biogeographic boundaries corresponding to Wallace’s and Lyddeker’s Lines are shown as thick black lines and define Wallacea as the region separating the continental Sunda shelf from Sahul. Coastlines are defined as −120 metres below present mean sea level, equivalent to the low sea level stand estimated at ~50ka.

Though most of these are endangered, all of them survive in the Sunda Shelf as living species today. In contrast, a wide variety of terrestrial animals larger than modern humans occupied the Sahul Shelf during the Late Pleistocene before the arrival of hominins in the region, including diprotodontid marsupial genera (*Diprotodon, Hulitherium, Maokopia*, and *Zygomaturus*) and large kangaroo genera (*Procoptodon, Simosthenurus*, and *Sthenurus*), along with the marsupial predator *Thylacoleo carnifex*, the gigantic monitor *Varanus priscus*, the large snake *Wonambi naracoortensis*, the land crocodiles *Quinkana* and *Pallimnarchus*, and the turtles *Meiolania* and *Ninjemys*. All of these species became extinct after human arrival and no native animal weighing > 60 kg occurs in Australia, Tasmania, or New Guinea today.

The disparate extinction histories of Sunda and Sahul may be partly explained by historical distribution of hominin species in these regions. While the long term presence of hominin species across the Sunda Shelf (including islands like Sumatra, Borneo, and Java)^1^ could have predisposed the local fauna to hominin impacts, the fauna across the Sahul shelf (including islands like New Guinea and Tasmania) were first exposed to hominins following AMH arrival. Extending this logic to the islands between Sunda and Sahul – i.e. Wallacea and the Philippines – might offer insights regarding their hominin occupation throughout the Pleistocene. Megafaunal survival shows markedly heterogeneous geographical distribution across these archipelagos – with Late Pleistocene extinction of *Rhinoceros* and *Bubalus* in the Philippines, and proboscidean species (*Elephas* and *Stegodon*) in the Philippines and Wallacea, matched by present-day survival of dwarf buffalo in the oceanic Philippines (Mindoro) and Sulawesi (*Bubalus* spp.), deer (*Rusa* spp.) and endemic pigs in the Philippines (*Sus* spp.) and on Sulawesi and its satellites (*Sus celebensis, Babirusa* spp.), and the Komodo dragon (*Varanus komodoensis*) on Flores and its satellites. Crucially, these patterns of megafaunal survival show a very close association with known areas of archaic human presence (from both fossil and stone tool records) in Flores, Sulawesi, and throughout the oceanic Philippines.

The most complete assemblages of living megafauna between Sunda and Sahul persist in central Sulawesi and on the island of Mindoro in the Philippines. Overall, this provides an intriguing indication that the best places to look for evidence for deep Pleistocene occupations where fossil hominin taxa have not yet been described may be on Sulawesi and its satellite islands, and in the oceanic Philippines south of Luzon. These may also be possible locations for the occurrence of Denisovans that are not yet evident from islands like Java, Flores, or Luzon, where hominin fossil records are now available. Conversely, the relatively large islands in the region that have no living megafauna, including Timor, Seram, or Halmahera, are unlikely to have had sustained hominin occupations prior to the arrival of modern humans. Together with the archaeological and genetic records, this approach suggests a broad region of archaic hominin presence in Wallacea (east of Wallace’s Line^33^ and west of Lydekker’s Line^34^ – Figure 3). In any case, our observations indicate that the first AMH populations to arrive in ISEA have most likely encountered a variety of hominin populations, no matter which route they took to enter Sahul^35–40^.

## Discussion

The lack of any detectable super-archaic introgression in our analyses stands in stark contrast to the strong evidence of Denisovan admixture with the ancestors of present-day ISEA populations^16–19,41,42^. Based on current paleoanthropological interpretations of *H. luzonenesis* and *H. floresiensis* as descendants of super-archaic hominin groups, our results indicate that interbreeding between these groups and AMH did not occur (at least at detectable levels), or that these encounters did not produce viable progeny, or that the offspring were viable but that these lineages have since died out. Evidence for super-archaic introgression into the ancestors of the Altai Denisovans^23^ and, possibly, Andamanese populations^24–26^, suggests that viable reproduction may actually have been possible, though further evaluation of these hypotheses is not possible given the available data.

An alternative explanation is that *H. luzonensis* and *H. floresiensis* belong to a hominin clade that is considerably less divergent from AMH than is currently accepted, possibly being the late-surviving descendants of an earlier radiation of a Denisovan-like lineage across ISEA. This would imply that hominin occupation of Flores (1.2Ma)^22,43^ and the Philippines (700ka)^44^ was not continuous and that the ubiquitous Denisovan ancestry across ISEA, which includes signals of admixture from a Denisovan population that diverged from the Altai Denisovans ~280ka^16^, results from AMH admixture with one or both of these groups. Indeed, the patterning of Denisovan ancestry across ISEA is consistent with separate Denisovan introgression events in the Philippines^18^ and, potentially, in Flores^17,19^. Despite the complete lack of support for this scenario from current morphological interpretations^7,9–11,20–22^, it is possible that pronounced dwarfism and prolonged periods of endemic island evolution for *H. floresiensis* and *H. luzonensis* complicate morphological assessments and phylogenetic placement of these groups. While it would simplify the search for ‘southern Denisovans’ if they could be linked with the dwarfed hominins of Flores and Luzon, existing archaeological and morphological data contradict such a possibility. By exclusion, this points to the region lying between the Philippines and Flores as a more likely location for the sources of Denisovan-like DNA.

A major complication in resolving these questions is the sparse Denisovan fossil record – currently consisting of one phalanx, a mandible, several teeth and a couple of bone fragments – which makes meaningful morphological comparisons very difficult. Clearly, further resolution of hominin prehistory of ISEA will greatly benefit from direct fossil and archaeological evidence of Denisovan presence in the region, with the potential for proteomic studies to assist in resolving phylogenetic relationships where DNA is not recoverable. Additionally, the patterns of megafauna survival across ISEA revealed in our study point to the widespread presence of archaic hominins east of Wallace’s Line^33^. This hints that much of the Denisovan ancestry found in modern human populations in ISEA, New Guinea, and Australia may have been locally acquired, emphasizing the need for more archaeological and genetic research across this region in the future.

## Methods

### Samples

We examined 426 individuals from 10 distinct populations (Table S1), taking advantage of publicly available data from previous genomic studies, and a recent effort to sequence hundreds of Indonesian genomes through the Indonesian Genome Diversity Project (IGDP)^16^. For a description of data preparation (SNP calling, QC, phasing) see Jacobs et al.^16^.

### Searching for signals of super-archaic admixture into modern humans

We searched for signals of super-archaic introgression in genomic sequences of AMH populations across the world, with a particular focus on ISEA and New Guinea (descendants from early AMH migrations into the region). These specific signatures are expected to include the existence of genetic variants that are not observed in Africa, and which exhibit levels of linkage disequilibrium compatible with introgression events ~60-50 ka, similarly to observations for Neanderthal and Denisovan introgressed segments. However, we expect deep divergence times between extinct ISEA hominins (*H. luzonensis* and *H. floresiensis*) and *H. sapiens* if we consider the former are not part of the Denisovan/Neanderthal clade and are instead related to *H. erectus*, or represent additional *Homo* lineages that split from AMH ~2 Ma or earlier. Hence, the putative introgressed super-archaic regions are expected to be highly divergent to orthologous modern human genome sequences. Importantly, the absence of a genome sequence for the extinct ISEA hominin groups makes this inference far more complex than for Neanderthal or Denisovan introgression, for which reference genomes are available. Therefore, we searched for super-archaic introgression in the genomes of contemporary human populations around the world using a novel, highly powerful Hidden Markov Chain model implemented by Skov et al.^45^ (termed here HMM_Archaic_), which is agnostic to the genome sequence of the putative archaic source. The rationale behind this strategy is that introgressed regions of the genome are enriched for genetic variants not seen in populations which have not admixed with the putative archaic source. In this case, we used African populations as an outgroup and assumed that these African populations have not interbred with Neanderthals, Denisovans, or any super-archaic source. It should be noted that the class of methods to which HMM_Archaic_ belongs are only indicative of archaic introgression. These methods might be prone to false positive detection of introgressed fragments due to incomplete lineage sorting or balancing selection maintaining old genetic diversity at specific selected loci. The HMM_Archaic_ method infers archaic admixture using a sliding-window approach after controlling for genetic diversity existing in an outgroup (e.g. African populations). We applied the method across all individuals from each of the ten sampled populations, using as an outgroup all individuals belonging to every African population contained in our dataset. After this, we further excluded positions where the Altai Neanderthal and Altai Denisovan individuals are heterozygous. We set the initial parameters to run HMM_Archaic_ following the author’s implementation, specifically: states =[‘Human’, ‘Archaic’]; starting_probabilities = [0.98, 0.02]; transitions = [[0.9995,0.0005],[0.012,0.98]], emissions = [0.04, 0.1]. Importantly, the method can be applied to phased data, and hence extract putative introgressing haplotypes rather than unphased regions, allowing for downstream analysis that is more sensitive to the independent histories of homologous chromosomal regions. Hence, the model was trained and implemented on phased data, which was obtained as described in Jacobs et al.^16^. We used a 1,000bp sliding-window approach, as performed in the original implementation of the method^45^, as the small size of the sliding-windows across the genome allows a fine-scale resolution of even small introgressed fragments where other methods^27–29^ are likely to fail.

The HMM_Archaic_ method outputs a posterior probability of introgression for each 1,000 bp window along each chromosome copy of each individual sample. These are called either ‘Human’ or ‘Archaic’ blocks, with each archaic block having posterior support >0.5; however, as we wish to focus on high-confidence introgressed blocks, we decided to drop archaic blocks with posterior probability support <=0.95. Therefore, the archaic blocks we examined were all regions directly estimated from HMM_Archaic_ with posterior probability >0.95, with no further changes such as merging of the inferred archaic blocks.

### Identifying Denisovan and Neanderthal introgressed fragments

We first sought to detect genomic signals of Neanderthal and Denisovan introgression using the CP^27^ and HMM^28,29^introgression-detection methods described in Jacobs et al^16^. These methods use phased data and seek to define haplotype blocks that are introgressed from an evolutionary relative of a sampled archaic genome, by detecting regions with a high density of variants that are shared with the archaic genome but not observed in an African outgroup sample. All parameters and details of the method implementations are given in Jacobs et al^16^

### Obtaining residual_Archaic_ blocks

We then focused on regions inferred to be introgressed using HMM_Archaic_^45^, which contain the introgressed fragments from Neanderthals and Denisovans and, potentially, additional introgressed signals not captured by CP or HMM. By subtracting the introgressed regions inferred to be of Neanderthal or Denisovan origin from CP and HMM, we produced a residual HMM_Archaic_ signal (residual_Archaic_) of blocks not overlapping Neanderthal or Denisovan fragments inferred with the other two methods. Specifically, for overlapping fragments, we subtract the overlapping HMM_Archaic_-CP/HMM regions, while still retaining the non-overlapping regions (refer to Supplementary Figure S8 for an illustration). This approach is allied to the residual *S\** calculated in Jacobs et al^16^, but differs in using more accurate phased archaic calls from HMM_Archaic_ and in the detail of the residual_Archaic_ block calling process. Note that identified residual_Archaic_ blocks may be in close proximity to Denisovan or Neanderthal introgressed regions (as is the case in Supplementary Figure S8) and that these blocks are not suitable for some downstream analyses such as introgression time estimation based on introgressed block length, as they may correspond to subparts of larger introgressed blocks. We decided to adopt this strategy as there is potential for super archaic blocks, in case they are present, to segregate close to, or overlap with, Neanderthal and Denisovan fragments, given the potential for non-random segregation of archaic blocks within the genome. While in the current work we do not present the results for an alternative strategy of completely removing Neanderthal and Denisovan blocks to estimate residual_Archaic_, the findings are qualitatively similar to the ones presented here.

### Looking at patterns of variation within residual_Archaic_ blocks

In order to further disentangle the patterns seen in residual_Archaic_ blocks, we looked at mutation-motif patterns. We defined the mutation motifs as 0 (ancestral) and 1 (derived), and a combination of [X, D, N, H], where ‘X’ represents the allelic state of a particular individual within an introgressed block (which can also be thought of as the test population – i.e., Papuan, East ISEA, West ISEA, etc), ‘D’ represents Denisova, ‘N’ represents Neanderthal, and ‘H’ represents an individual from an African population (in our case Ju’hoan - SS6004473). While all African variation was removed from the dataset prior to running HMM_Archaic_ (as Africans form the required outgroup), we reintroduced SS6004473 variation subsequently and for this specific analysis only. This means that, for example, the mutation motif [1001] is seen when X shares the derived allele with the African individual, and Neanderthal and Denisovan are ancestral; likewise, the mutation motif [1000] indicates regions where X carries a derived allele that is not observed in the African individual, Neanderthal or Denisovan. Hence, in the case of super-archaic introgression into modern humans, an enrichment in [1000] and [0111] motifs within introgressed blocks is to be expected.

### Variation in motif proportion as a function of physical distance to introgressed regions

We investigated the proportion of different motifs as a function of physical distance to the putatively introgressed regions. In this case we divided the analyses into patterns seen within all HMM_Archaic_ introgressed fragments and those seen residual_Archaic_ fragments (Supplementary Figure S7). In this analysis, we define mutation motifs as [X, D, N, Af] where a single human outgroup is now represented by an indicator Af, 1 indicates that a variant is found in the derived state in one or more individuals in the African outgroup, and 0 indicates that the derived state is not observed. Thus, we are specifically focusing on whether variation is found at all in an African sample rather than a single African individual. When all HMM_Archaic_ fragments within the Papuan population are considered, we observe an excess of [1100] and [1010] motifs, compatible with introgression from Denisovan and Neanderthal into Papuan genomes, respectively, along with a sharp decrease of [1001] (where X shares a derived allele with Africa) motifs. These signatures consistently indicate Neanderthal and Denisovan introgression into Papuan genomes. When considering residual_Archaic_ fragments only, we observe a sharp increase in the [1000] motif (as expected) coupled with a reduction in the [1100] and the [1010] motifs (signals of Denisovan and Neanderthal introgression, respectively), suggesting that remaining fragments do not show a clear signal of known archaic introgression. These Neanderthal and Denisovan signals increase in the regions around residual_Archaic_ blocks, indicating that they are often nested within introgressed Neanderthal and Denisovan sequences. This is an important observation, suggesting that much of the signal is contributed through known introgression, in support of the absolute increase in residual_Archaic_ in Papuan populations. Indeed, the definition of residual_Archaic_ does not exclude the detection of regions showing coalescent histories consistent with super-archaic introgression from within Denisovan and Neanderthal introgression (as would likely be the case for the blocks shown in example schematic Supplementary Figure S8), and variation in the coalescent histories within blocks sharing the same introgression source is likely. While this suggests that residual_Archaic_ blocks may be retrieving super-archaic signals from within Denisovan and Neanderthal introgressing populations, we suggest that more data and more focused analysis, beyond the scope of this paper, are necessary to assess the significance of these patterns. The sharp decrease in the [1001] motif observed in all HMM_Archaic_ blocks is replaced by a peak in residual_Archaic_ blocks, and a slight increase in the [0111] motif is now visible. In both cases, these indicate deep coalescence of residual_Archaic_ blocks not associated with the sampled Neanderthal or Denisovan sequences. While the [0111] signal is of particular interest in the context of super-archaic introgression, the lack of any global peaks in this motif (Figure S2) and elevated [1100] and [1010] signals surrounding residual_Archaic_ blocks argues that it more likely reflects deep coalescent histories within Denisovan and Neanderthal introgressed blocks than super-archaic introgression.

### Motif proportion differences are correlated with known archaic ancestry

We explicitly test for a correlation between Neanderthal and Denisovan ancestry and motif proportions within residual_Archaic_ blocks between populations. Supplementary Figures S3 and S4 show the correlation between inferred Denisovan, and Neanderthal ancestry, respectively, and the proportion of different motifs, across all individuals. Interestingly, we find both positive and negative correlations between the proportion of different motifs and the detected amount of Denisovan and Neanderthal ancestry. In fact, these correlations are statistically significant for all but two motifs when regressing on Denisovan ancestry, [0100] (P-value 0.289) and [1110] (P-value 0.618), and for all but one motif when regressing on Neanderthal ancestry, [1110] (P-value 0.221). These results are in agreement with the observations from simulations with no *super-archaic* introgression, which show that residual_Archaic_ sequence is essentially dominated by introgressed Neanderthal and Denisovan fragments that are undetected by both HMM and CP.

### Simulating super-archaic introgression using msprime

In order to test the power of our experimental design to detect introgression from a highly diverged human lineage into the ancestors of ISEA populations/Australo-Papuans, we implemented a series of neutral coalescent simulations using the software *msprime^31^*. The simulations use demographic parameters derived from Malaspinas et al.^32^, which models Aboriginal Australian history from full genome data from modern Australian and Papuan populations. The structure and parameters describing the standard demography (i.e. excluding possible super-archaic introgression) followed the maximum likelihood model output (V. Sousa, pers. comms.). Briefly, we simulated a total of 35 African and 30 Australian individuals, and one Altai Denisovan individual that split from human populations 20,255 generations prior to the present, while African and Australian populations split from one another 3,916 generations ago. Additionally, we included one super-archaic individual, that splits from the Human-Neanderthal-Denisova clade 70,000 generations in the past, to mimic the deep split assumed for *H. floresiensis* and *H. luzonensis*, with haploid Ne = 13,249. Following Malaspinas et al.^32^, Neanderthal (2.4%) and Denisovan (4.0%) introgression events were simulated at, respectively, 1,853 and 1,353 generations in the past, with the introgressing lineages being related to the Altai individuals, and additional minor Neanderthal contributions to the Eurasian clade (1.1%) and Australian clade (0.2%) at 1566 and 883 generations ago, respectively. For the super-archaic admixture, we assumed an introgression event occurring 1,353 generations ago. We set the mutation rate to 1.4e-8/bp/generation and the recombination rate to 1e-8/bp/generation and simulated, per individual, a total of 300 chromosomes of 10Mb in length each. This strategy allowed us to obtain a total simulated sequence that roughly matches the size of the human genome for each individual (~3Gb of sequence), while ensuring sufficient independent replication. Importantly, after running the simulations, we sampled 65 human individuals (35 African and 30 Australian genomes), an Altai Neanderthal and an Altai Denisovan (related to, respectively, the introgressing Neanderthal and Denisovan populations), and one super-archaic individual.

A major advantage of using *msprime* to implement coalescent simulations is that the software allows the genealogy of each portion of simulated sequence to be traced back through time, including the migration of genomic regions between archaic and human populations (i.e. introgression). This means that, for each individual, we are able to know the exact amount and location of the introgressed segments, and are thus able to directly compute the strength of our approaches for detecting super-archaic introgression in the empirical data.

### Models of super-archaic introgression

We initially implemented two models of super-archaic introgression: a model containing 2% introgression into the ancestors of Australians occurring at the same time as Denisovan introgression, and a second model without super-archaic introgression (0%). To estimate the power of our analytical framework to detected super-archaic introgression at low levels of admixture, for each simulated individual we created datasets with ~1% and ~0.1% super-archaic introgression by masking a specific proportion of super-archaic blocks in the 2% model. Specifically, this was achieved by 1) randomly sampling a proportion of introgressed super-archaic regions in each individual; and 2) merging all the regions sampled across all individuals and masking these merged super-archaic regions across all simulated individuals. This strategy ensured that the masked super-archaic regions were the same across all individuals. We were able to reduce the amount of super-archaic ancestry present in the simulated sequences to ~1% and ~0.1% by randomly sampling, per individual, ~10% and ~50% of introgressed super-archaic regions, respectively. Due to the masking of the introgressed regions, the 1% and 0.1% models contained slightly less genetic sequence than the 0% an 1% models (~2.88Gb and ~2.65Gb simulated sequence, respectively); however the masking did not alter the average proportion of introgressed sequences observed from either the Denisovan of Neanderthal lineages (Supplementary Figure S12).

### Power to uncover archaic introgression

We evaluated the performance of the analytical pipeline by comparing the results from our empirical data to four models of Australian-super-archaic admixture at different introgression levels (i.e. 2%, 1%, 0.1% and 0%). First, we estimated the power of each of the three detection methods utilized to compute archaic introgression in the empirical data; i.e. CP, HMM and HMM_Archaic_. Analogous to the implementation in the empirical data, before running HMM_Archaic_, we excluded all variation present in the 35 simulated African genomes, along with positions for which the Altai Neanderthal and Denisovan individuals were heterozygous. Supplementary Figure S9 shows the True Positive Rate (TPR) of each method to detect archaic introgression. The TPRs were estimated as the length of detected regions that overlap the simulated introgressed regions over the total length of simulated introgressed regions (in base-pairs). It was possible to estimate the TPR separately for introgression from the Neanderthal and Denisovan lineages for CP and HMM, though not for HMM_Archaic_ (which does not require a reference).

Both CP and HMM consistently detect Neanderthal introgression at a higher rate than Denisovan introgression, irrespective of the amount of super-archaic introgression present in the simulations (Supplementary Figure S9). Considering that both CP and HMM rely on the availability of a reference sequence for the putatively introgressing archaic population, this observation is consistent with the fact that the simulated introgressing Neanderthal population is genetically closer to the reference Altai Neanderthal than the simulated introgressing Denisovan population is to the reference Altai Denisovan. Nevertheless, both methods seem to perform only slightly better in the absence of super-archaic introgression, presumably because, at least in the case of CP, a very small proportion of inferred Neanderthal and Denisovan introgression derives from super-archaic introgression (see below). HMM_Archaic_ has extremely high power to detect super-archaic segments (Supplementary Figure S9, top left) and, even though power decreases at lower levels of super-archaic introgression, it is always higher than the detection power for Neanderthal or Denisovan introgression across all four models (Supplementary Figure S9).

### False positive rate

We next examined the False Positive Rate (FPR) of each method to detect archaic introgression. For the CP and HMM methods we define FPR as the proportion of sequence misassigned to a particular archaic population when that sequence is either introgressed from another hominin lineage or is from the human genealogy. For HMM_Archaic_ we simply estimated the proportion of sequence misassigned as archaic that overlaps simulated human regions. The results are shown in Supplementary Figures S10 and S11. Both CP and HMM have relatively high FPRs when inferring Neanderthal introgression that actually results from Denisovan introgression (~40%), and vice-versa (~35%) (Supplementary Figure S11). As expected, given the closer relationship of the introgressing Neanderthal population to the reference Altai Neanderthal compared to the introgressing Denisovan population to the reference Altai Denisovan, the FPR for CP and HMM is higher for Denisovan segments that were missasigned as Neanderthal (Supplementary Figure S11, right panel) than vice-versa (Supplementary Figure S11, left panel). This pattern is also consistent with a close genetic relationship between Neanderthals and Denisovans, and the persistence of shared ancestral genetic diversity between the two species (incomplete lineage sorting). Importantly, however, the FPR of both methods is extremely low when inferring Neanderthal or Denisovan introgression when it either did not occur (Supplementary Figure S11, middle columns – ‘Human’) or the source was super-archaic (Supplementary Figure S11, left columns – ‘super-archaic’). Hence, our simulation results demonstrate that a negligible amount of introgressed super-archaic sequence will be mistaken for Neanderthal or Denisovan introgression by CP and HMM. Finally, our stringent approach for detecting archaic introgression using HMM_Archaic_ (posterior probability >0.95, see above) results, as expected, in virtually no false positives in the simulations (Supplementary Figure S10) – i.e. a negligible portion of archaic HMM_Archaic_ overlaps with human genealogies.

### Estimation of residual_Archaic_

We next investigated how this combination of TPRs and FPRs translated into the actual amount of recovered sequence. The results are shown in Supplementary Figure S12, contrasting the total amount of simulated introgression versus the total amount detected for each archaic species using the different methods. Notably, the amount of Neanderthal and Denisovan introgression detected by HMM_Archaic_ consistently increases as the amount of super-archaic ancestry declines (see below – *Effects of super-archaic ancestry to detect Neanderthal/Denisovan introgression*). In contrast, the amount of Neanderthal and Denisovan detected by both CP and HMM is essentially independent from the amount of super-archaic ancestry present (as expected from the TPRs shown in Supplementary Figure S9). As described above, the masking strategy adopted to reduce the amount of super-archaic in the simulations meant that models 1% and 0.1% contain a reduced amount of introgressed Neanderthal and Denisovan sequence overall (see explanation in *Power to uncover archaic introgression).Therefore*, we also present a corrected amount of simulated and detected archaic sequences by normalizing the total amounts to match the total amount of sequence considered in the empirical data (Supplementary Figure S12, panel b). This strategy also allowed us to compare the simulations directly to the results obtained for the empirical data, namely in terms of total residual_Archaic_ sequence present. After determining the total detected sequence in each method, we obtained the residual_Archaic_ regions by removing those regions that overlap with either the CP or HMM detected blocks (residual_Archaic_ in Figure 2, overlapping blocks shown as overlapArchaic).

### Effects of super-archaic ancestry to detect Neanderthal/Denisovan introgression

An interesting picture emerges when we consider the behaviour of HMM_Archaic_ in the presence of super-archaic introgression. The ability of HMM_Archaic_ to detect Neanderthal and Denisovan introgression is severely depleted at higher levels of super-archaic introgression, which appears to dominate the amount of detected archaic ancestry: less than 25% of truly introgressed Neanderthal and Denisovan sequences were detected when we simulate 2% super-archaic introgression, versus ~40-60% true rates for a model containing 0% super-archaic introgression (Figure S9, top panel). This pattern is consistent with the power of HMM_Archaic_ being proportionate to the divergence between the introgressing archaic population and the outgroup human population (i.e. Africa). Importantly, we have simulated a super-archaic source whose divergence to modern humans is significantly higher than that of Neanderthals and Denisovans to mimic introgression from *H. floresiensis* and *H. luzonensis*, assuming that the latter are earlier diverging lineages of *Homo* (see Methods). There is a considerably higher agreement between HMM_Archaic_ and both CP and HMM for a model with no super-archaic introgression compared to a model containing even 0.1% super-archaic introgression (Figure 2). The most important signal for differentiating these scenarios, which have similar total simulated residual_Archaic_, is the concordance between HMM_Archaic_ and CP/HMM. Specifically, the excess divergence of super-archaic introgressed sequences means these blocks contain a higher amount of non-African variants and, therefore, are more efficiently detected by HMM_Archaic_. However, this process simultaneously impacts the internal optimisation of HMM_Archaic_ emission parameters, causing the algorithm to seek more divergent introgressed blocks, which reduces the TPR for detecting known Denisovan and Neanderthal blocks. This is consistent with HMM_Archaic_ having a higher TPR for introgressed Neanderthal and Denisovan sequences when no super-archaic introgression is present, which in turn leads to a higher amount of Neanderthal and Denisovan sequence detected by all three methods (Figure 2). This behaviour causes the concordance between methods to drop, and the residual_Archaic_ signal to increase as a proportion of total HMM_Archaic_, even when simulating minimal amounts of super-archaic introgression. The higher concordance between HMM_Archaic_, CP, and HMM for the 0% model translates into a 27% proportion of residual_Archaic_ in this model (Figure 2c) – consistent with residual_Archaic_ regions computed in the empirical data (between ~15% in Papuan genomes and ~22% in West Eurasian genomes – Supplementary Figure S1) – and in contrast to ~33% to 60% for models with >=0.1% super-archaic introgression. Importantly, in simulations containing higher proportions of super-archaic ancestry (1% and 2% models), we observe a much higher proportion of residual_Archaic_ sequence.

### Investigating mutation motifs within residual_Archaic_ simulated models

In order to further investigate the nature of genetic diversity within residual_Archaic_ regions, we performed similar mutation motif analyses to those used in the empirical data (see above). In particular, we investigated the amount of shared ancestral and derived alleles between individuals carrying the residual sequence (i.e. test population), the simulated Altai Denisovan, the simulated Altai Neanderthal, and a simulated African genome – again, while all African variation was excluded from HMM_Archaic_ analyses, we randomly sampled one individual and investigated allele sharing within residual_Archaic_ regions after running the method.

## Acknowledgements

We thank Christian Huber and Joshua Schmidt for useful discussions on the genetic analyses, and Kieren Mitchell and Fernando Racimo for comments on the manuscript. This work was supported by ARC Indigenous Discovery Grant IN180100017 (JCT and RT) and ARC Laureate Fellowships FL100100195 (CMST) and FL140100260 (AC). CS acknowledges funding from the Calleva Foundation and The Human Origins Research Fund.

## Author contributions

JCT, KMH, GSJ, MPC, GH and JT designed the methods and undertook the analyses. JCT, KMH, RT, GSJ, AC, CS, CMST and MPC wrote the manuscript with input from all authors.

