## Supplementary material for "Introgression, hominin dispersal and megafaunal survival in Late Pleistocene Island Southeast Asia": Supp_Info

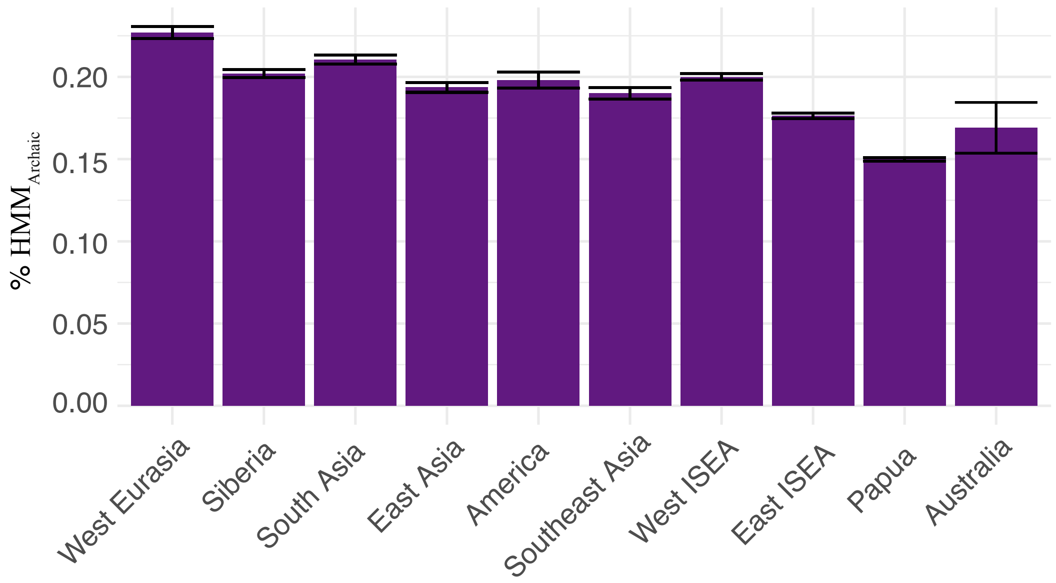


**Fig. S1.** Barplots showing the mean proportion of $residual_{Archaic}$ in real data. Proportions were obtained as the total amount of $HMM_{Archaic}$ sequence with probability of being archaic >0.95 and that were not detected by CP and HMM as Neanderthal nor Denisovan sequence. Mean $residual_{Archaic}$ was obtained across all individuals in different populations. Error bars denote the standard error on the mean values.

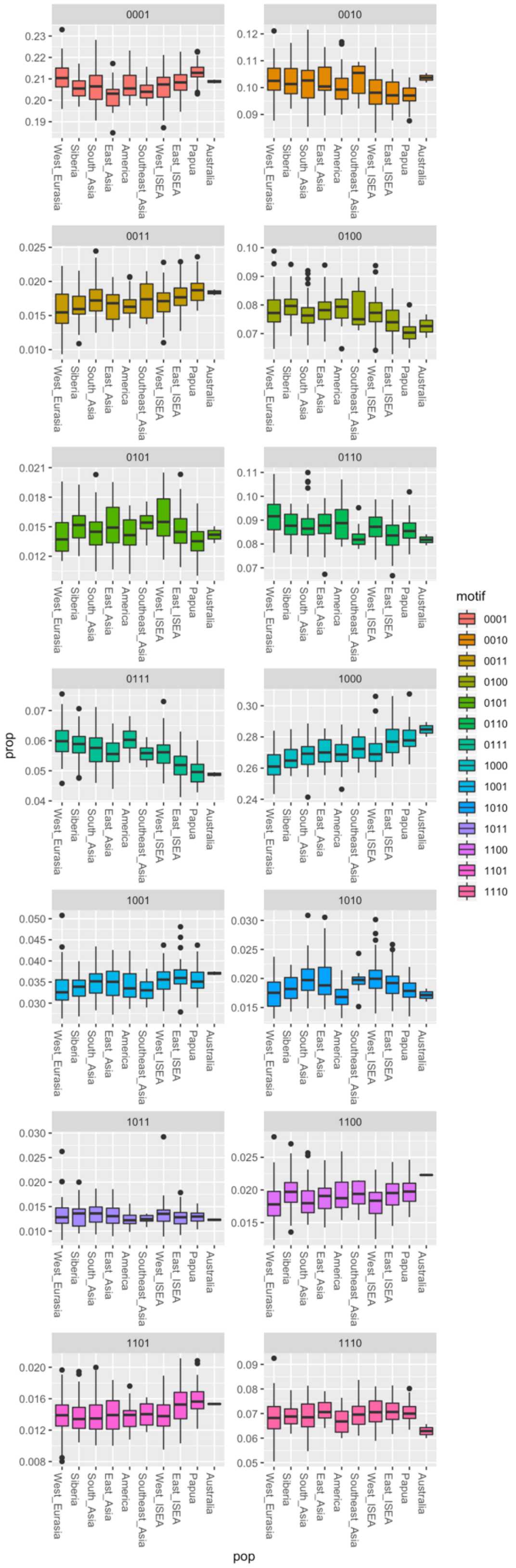


**Fig. S2**. Boxplots of the proportion of mutation motifs of the type [X, D, N, H] within residual$_{\text{Archaic}}$ blocks in different populations. A total of 14 mutation motifs were considered (we excluded motifs 0000 and 1111, where all individuals carry the ancestral or derived alleles, respectively). X represents an individual from a particular population, D represents Denisovan, N represents Neanderthal and H represents Human. The allelic states 0 and 1 correspond, respectively, to the ancestral and derived allelic states. Instead of considering each population value, here we consider the proportion of different motifs in each individual and plot the distribution of these proportions in each population, i.e. considering the motif proportions across individuals in different populations.

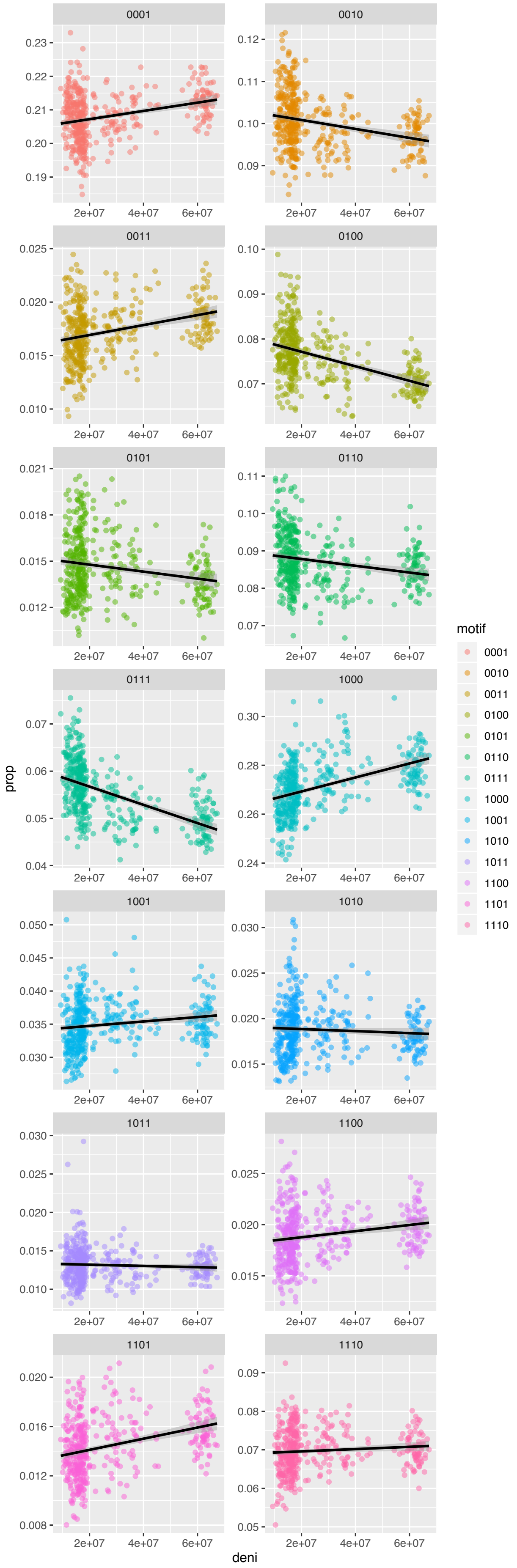


**Fig. S3.** Proportion of different mutation motifs of the type [X, D, N, H] (y-axis) and Denisovan ancestry (x-axis) within residual$_{\text{Archaic}}$ blocks across all individuals (represented by filled circles). Denisovan ancestry is represented as the total amount (Mb) of inferred Denisovan introgressed segments in each individual.

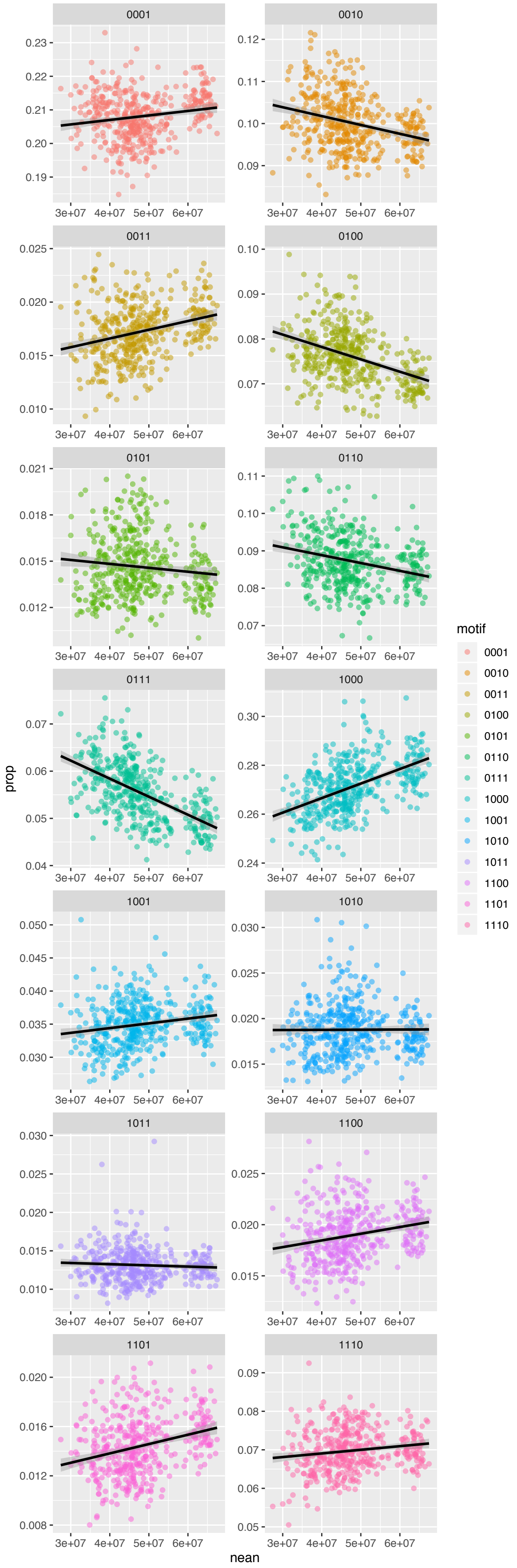


**Fig. S4.** Proportion of different mutation motifs of the type [X, D, N, H] (y-axis) and Neanderthal ancestry (x-axis) within residual$_{Archaic}$ blocks. Similar to Figure S11, Neanderthal ancestry is represented as the total amount (Mb) of inferred Neanderthal introgressed segments in each individual.

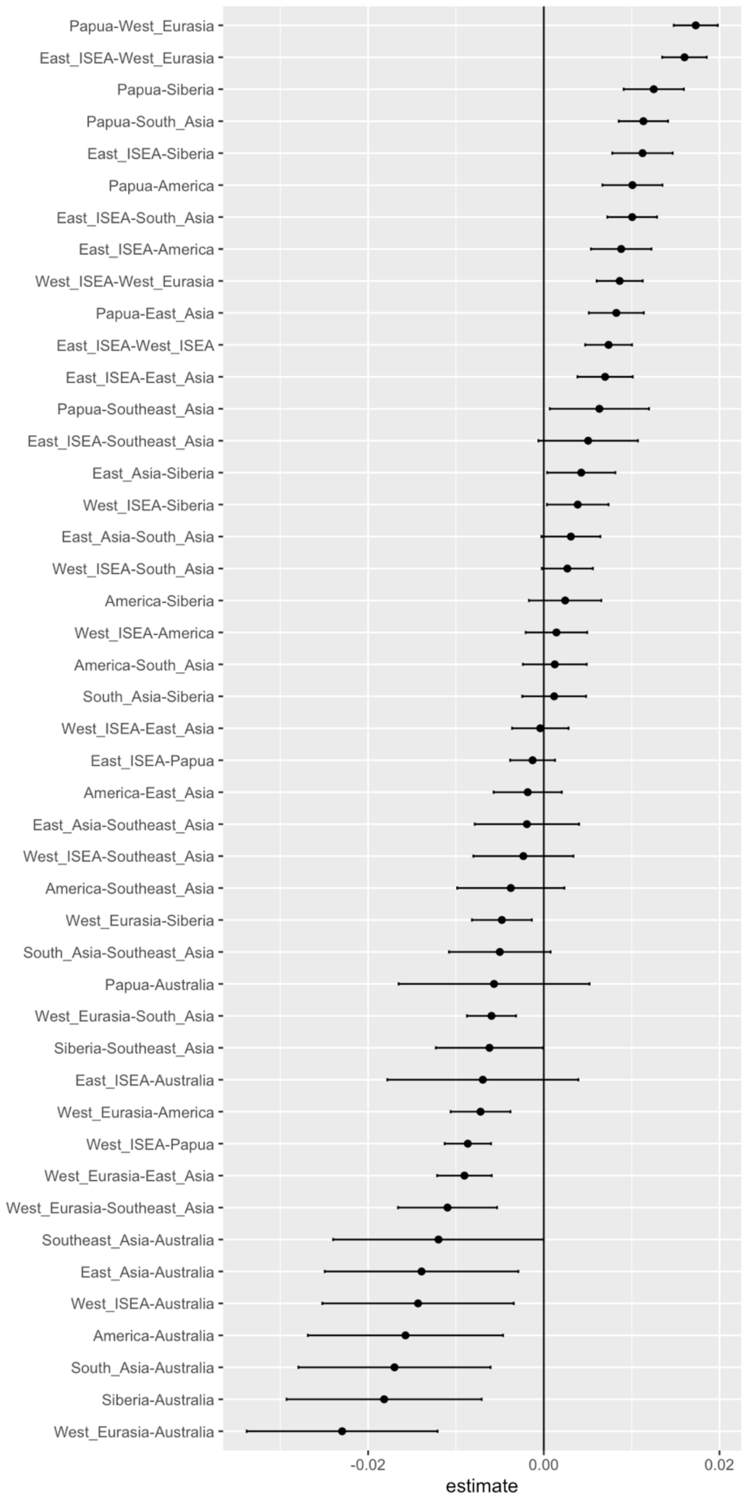


**Fig. S5.** Differences in the proportion of 1000 motifs (of the type [X, D, N, H]) between different populations. The error bars are 95% confidence interval for the true difference in proportions between the given populations. These confidence intervals were obtained using simultaneous inference procedures[1].

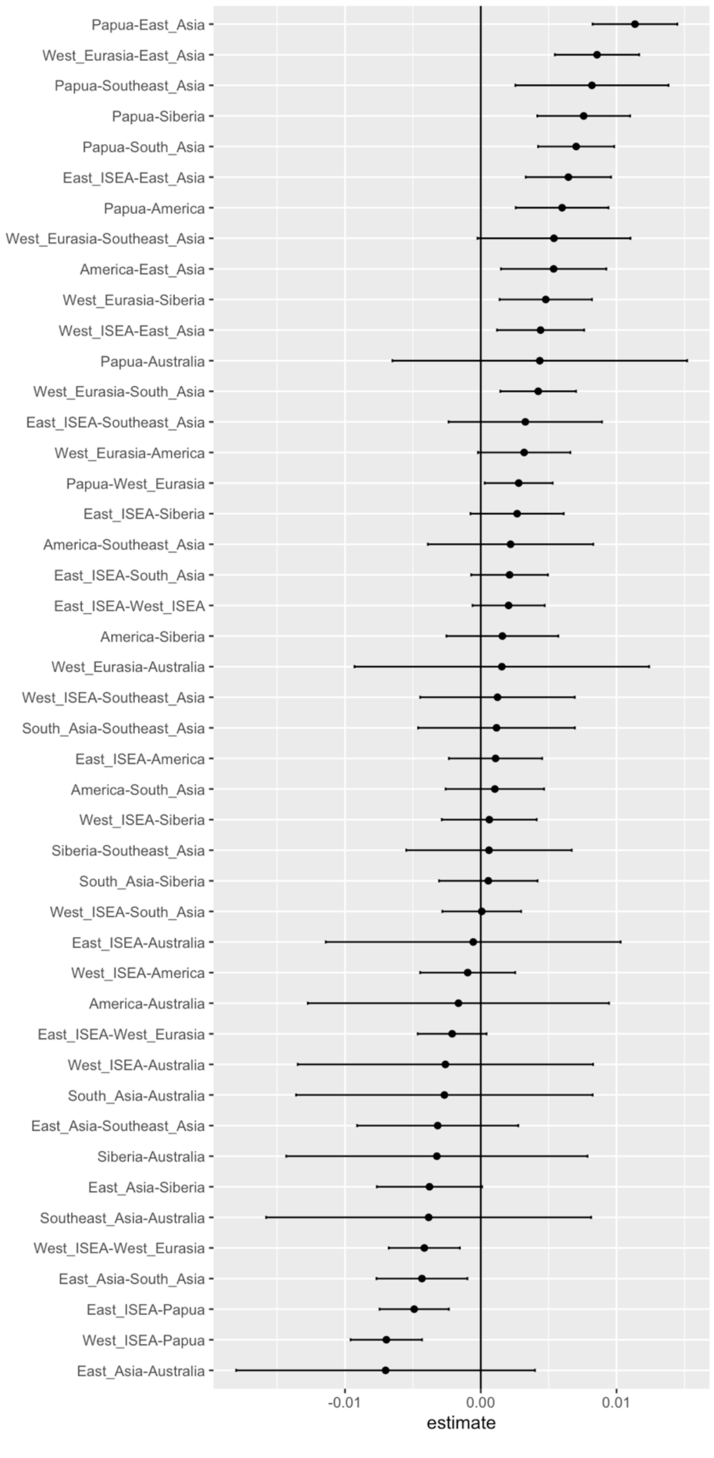


**Fig. S6.** Differences in the proportion of 0111 motifs (of the type [X, D, N, H]) between different populations. The error bars are 95% confidence interval for the true difference in proportions between the given populations. These confidence intervals were obtained using simultaneous inference procedures[1].

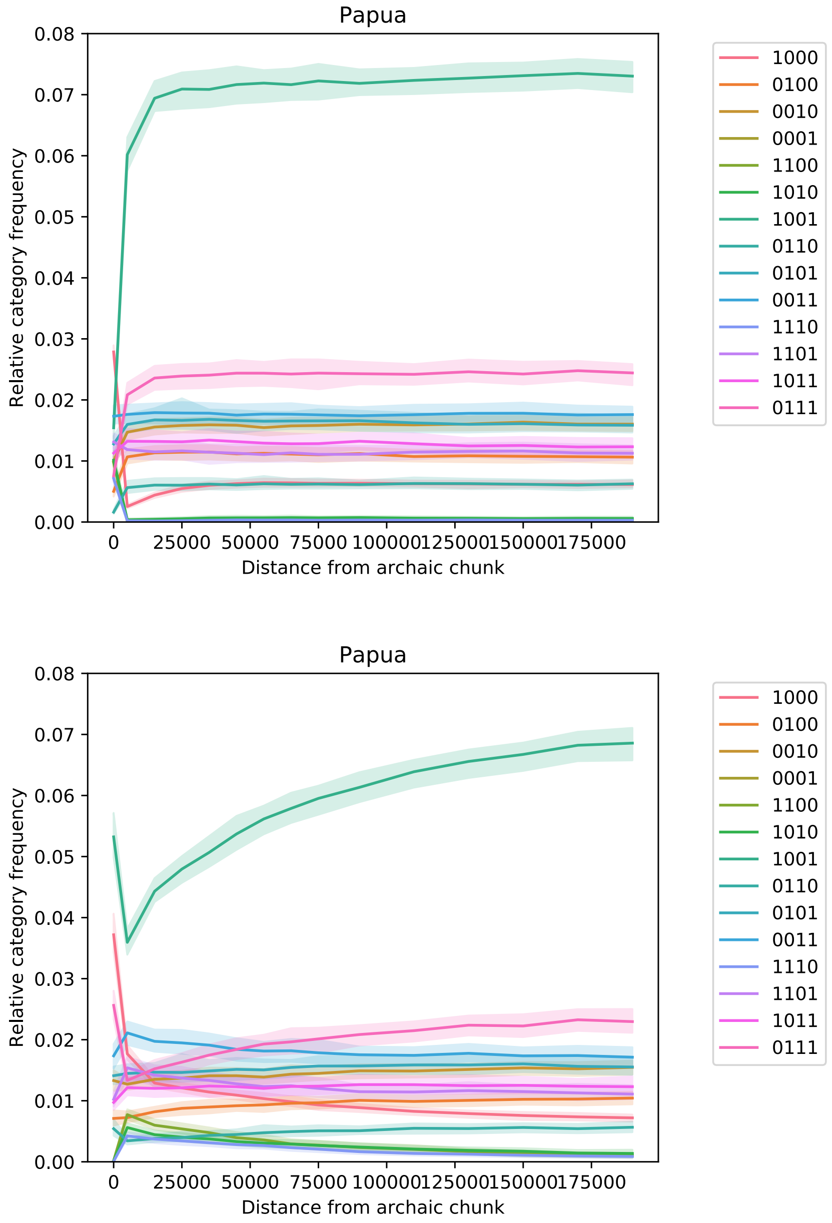


**Fig. S7.** Proportion of different mutation motifs of the type [X, D, N, H] as a function of spatial distance to $HMM_{Archaic}$ blocks. Top panel: all $HMM_{Archaic}$ blocks. Bottom panel: $residual_{Archaic}$ blocks. Types are as defined in Figure S2. Please note that distances to $HMM_{Archaic}$ blocks shown in the X-axis are calculated in base pairs.

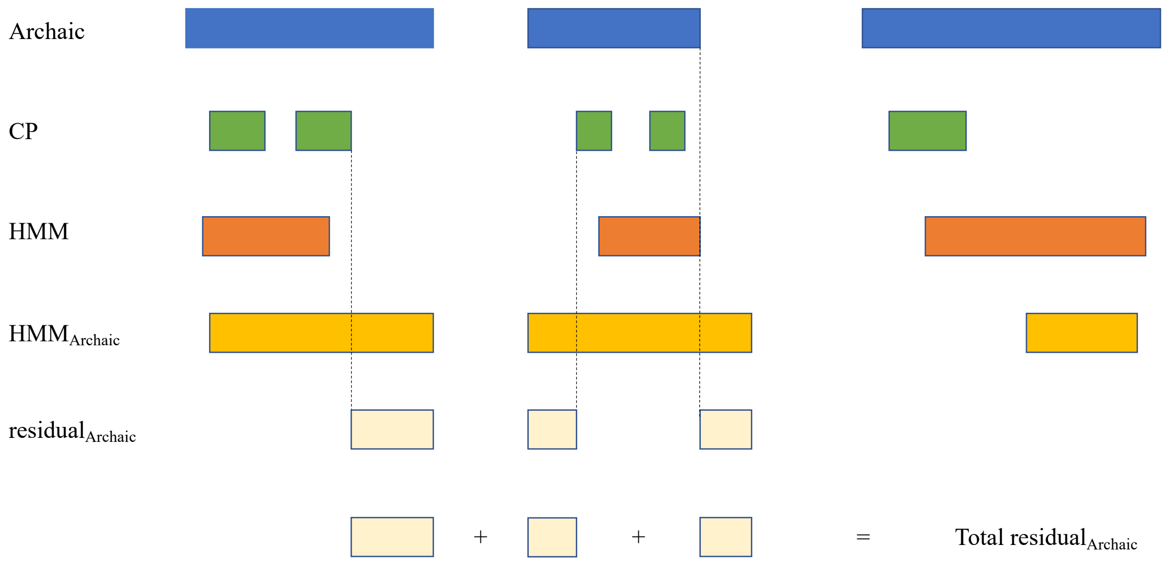


**Fig. S8.** Schematic representation on the calculation of residual$_{Archaic}$ blocks in a particular individual. Blue rectangles are meant to represent true archaic introgressed fragments in a particular individual. Coloured rectangles below represent the hypothetical detection of introgressed regions using different methods: CP (green), HMM (orange) and HMM$_{Archaic}$ (yellow). residual$_{Archaic}$ is represented in the rectangles at the bottom (light yellow), as the fraction of HMM$_{Archaic}$ that does not overlap with CP and HMM.

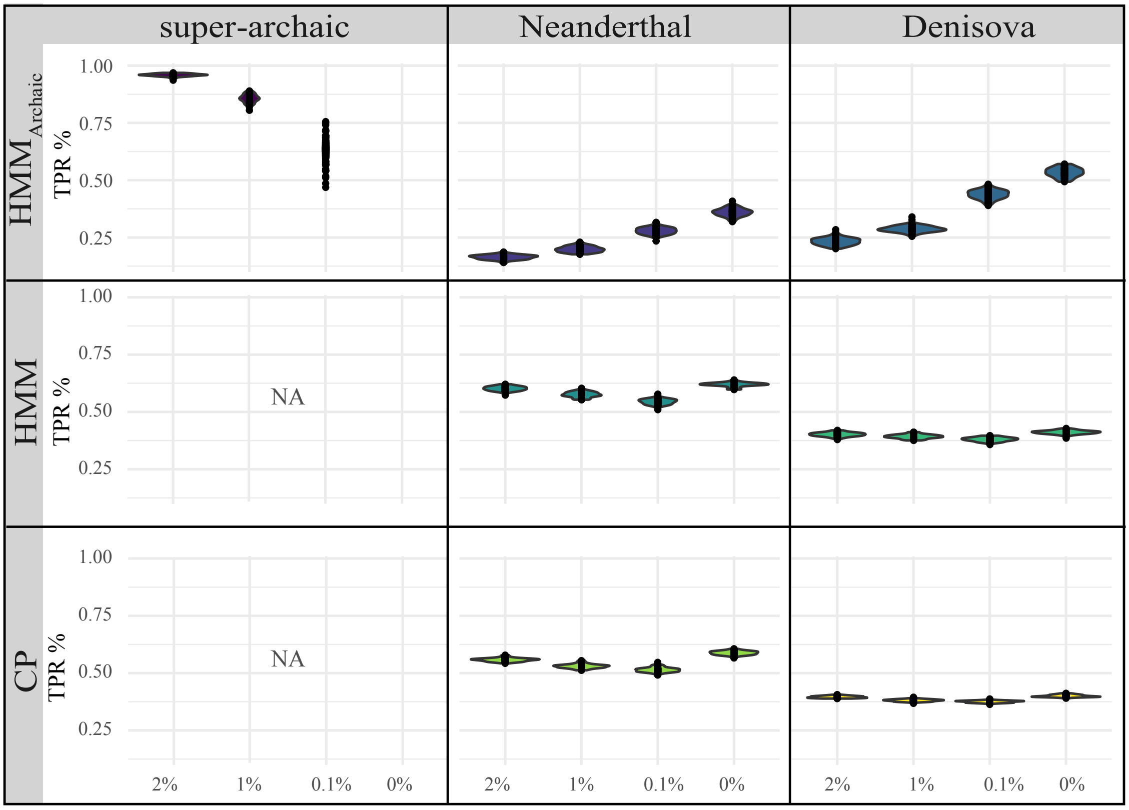


**Fig. S9.** True positive rate (TPR) of CP, HMM and $HMM_{Archaic}$ to detect introgressed regions in simulated data. TPRs were computed by dividing true positives over the total length of simulated introgressed sequence in base pairs. TPRs were calculated for different archaic sources (represented in separate panels) and simulated models of super-archaic introgression (x-axis).

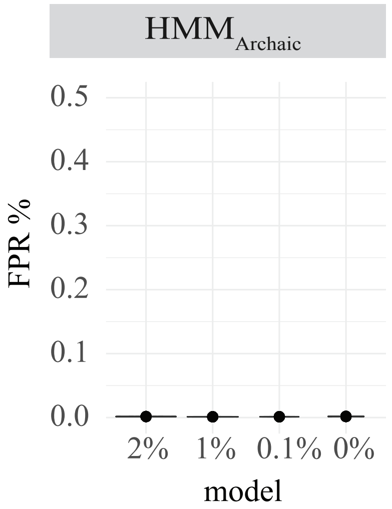


**Fig. S10.** False positive rate (FPR) of $HMM_{Archaic}$ when considering introgressed fragments with probability >0.95 across the different simulated models of super-archaic introgression (x-axis). Only fragments inferred as archaic (P>0.95) overlapping 'human' simulated genealogies are considered.

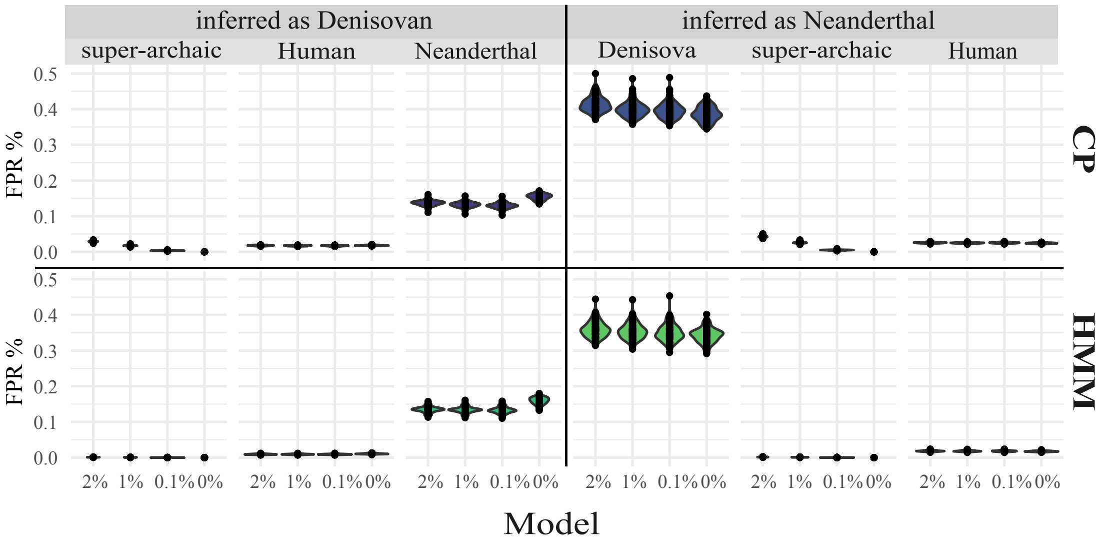


**Fig. S11.** False positive rate (FPR) of CP and HMM to detect introgressed Neanderthal and Denisovan regions in simulated data. FPRs were computed by dividing false positives over the total length of simulated introgressed sequence in basepairs. FPR is shown for all inferred blocks as either Neanderthal or Denisovan when these blocks in fact overlap introgressed regions from a super-archaic source, Human, and Denisovan or Neanderthal, respectively. FPRs were calculated separately for each simulated model (x-axis).

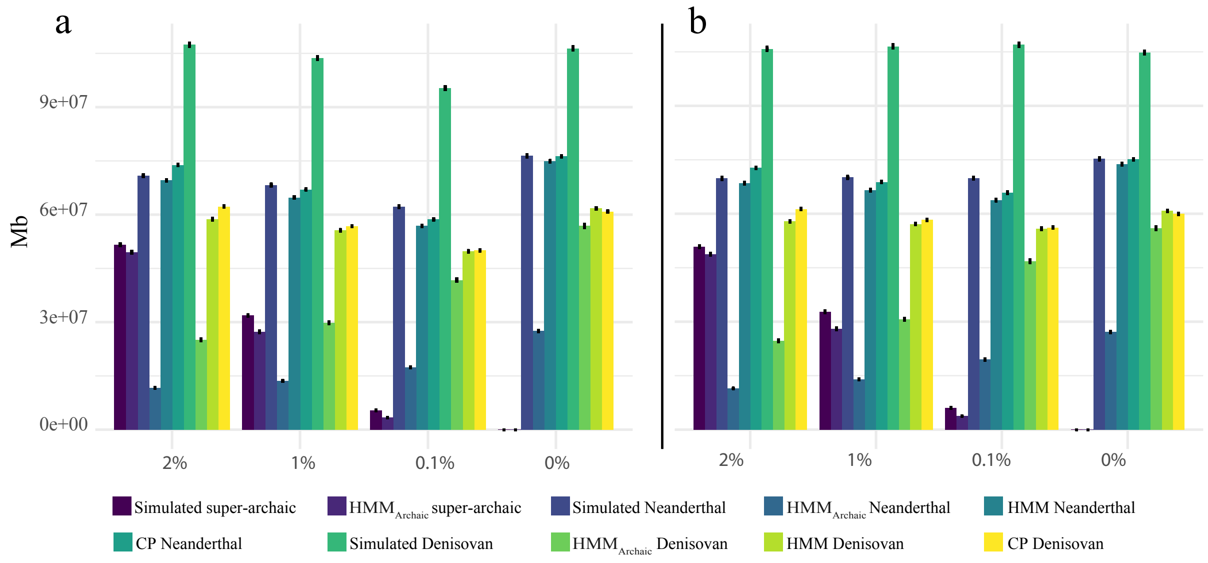


**Fig. S12.** Total amount of simulated super-archaic, Neanderthal and Denisovan sequence (in Mb), and respective amount of detected archaic sequence for each of the three methods. a) Shows the actual simulated sequence in each method (note that the amount of simulated Neanderthal and Denisovan also decreases in models 1% and 0.1% after masking super-archaic sequences, as some super-archaic segments in particular individuals overlap Neanderthal and Denisovan segments in other individuals). b) Corrected amount of simulated sequence by normalizing the proportion of introgressed segments by the actual amount of considered genome in the real data (note that unlike what is observed in panel A, the amount of Denisovan and Neanderthal, after correction, in models 1% and 0.1% is now very similar to models 2% and 0%).

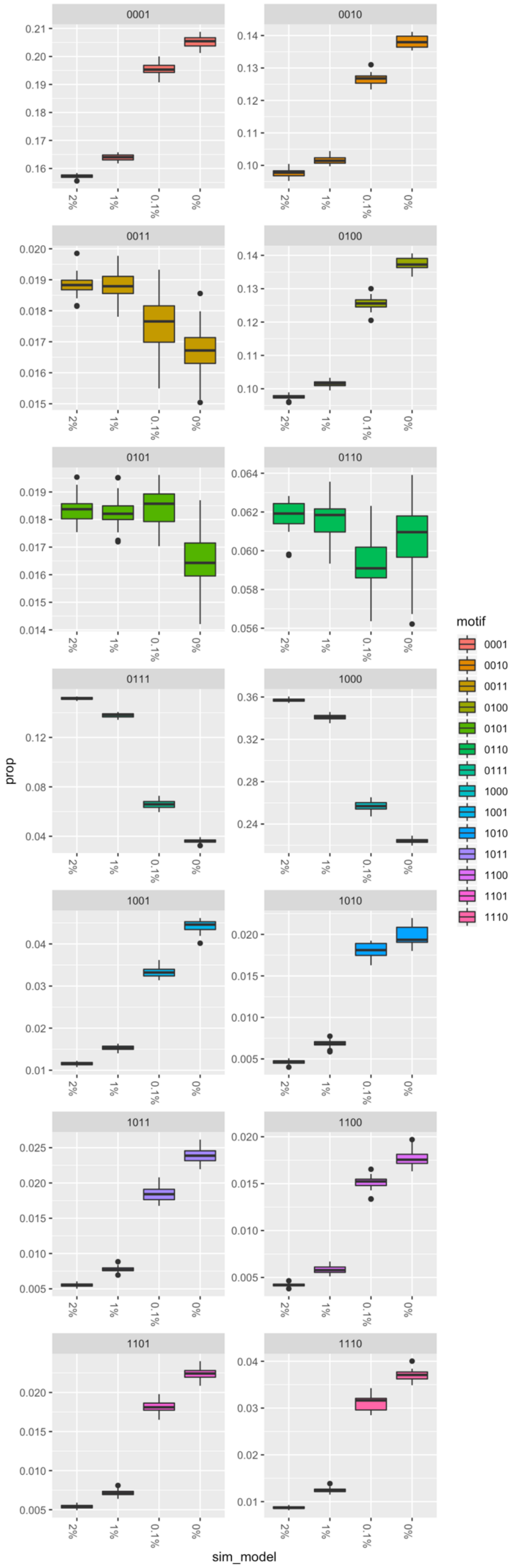


**Fig. S13.** Proportion of different mutation motifs of the type [X, D, N, H] (y-axis) within residual$_{\text{Archaic}}$ blocks in different simulated models of super-archaic introgression (x-axis). As above, only 14 mutation motifs were considered (i.e. we excluded motifs 0000 and 1111, containing sites where all individuals carry the ancestral or derived alleles, respectively).

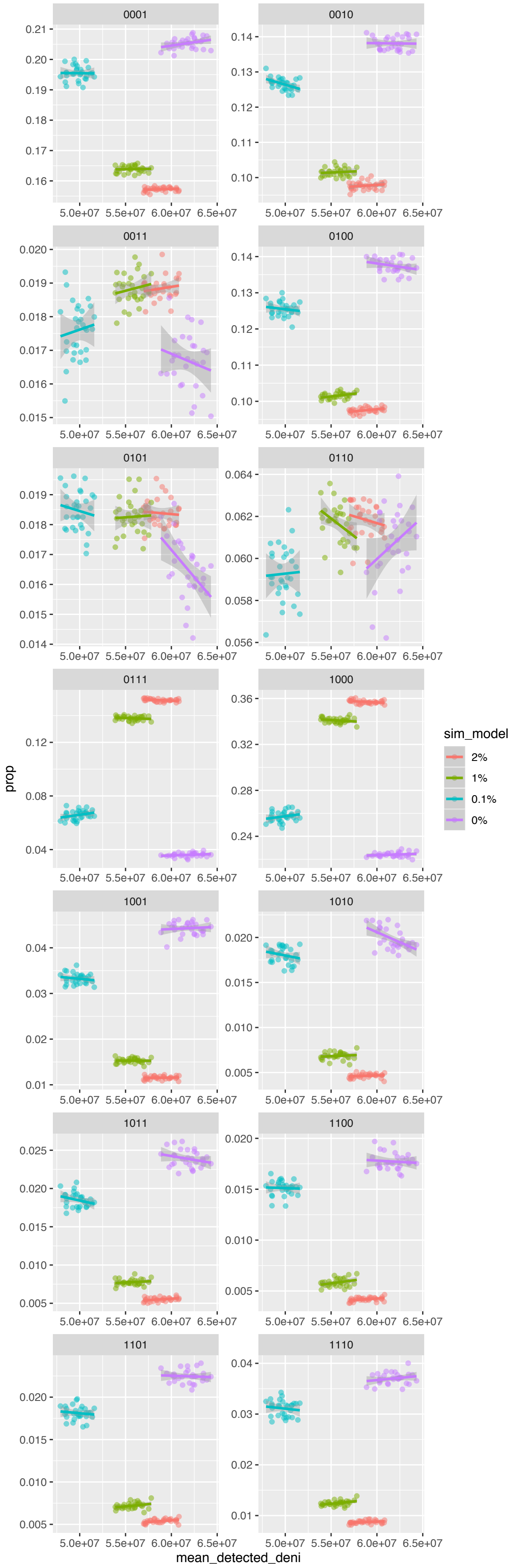


**Fig. S14.** Correlation between detected Denisovan ancestry and different mutation motifs of the type [X, D, N, H] in simulated data. The proportion of mutation motifs (y-axis) was calculated for each simulated individual in different models of super-archaic introgression. The total amount of Denisovan is shown in basepairs (x-axis) as detected by *HMM* in the different models. Note that models with no super-archaic introgression always resulted in a higher total detected amount of Denisovan introgression, in similar amounts to the real data. Once again, we disregarded mutation motifs of the types 0000 and 1111 where all individuals carry the ancestral or derived alleles, respectively.

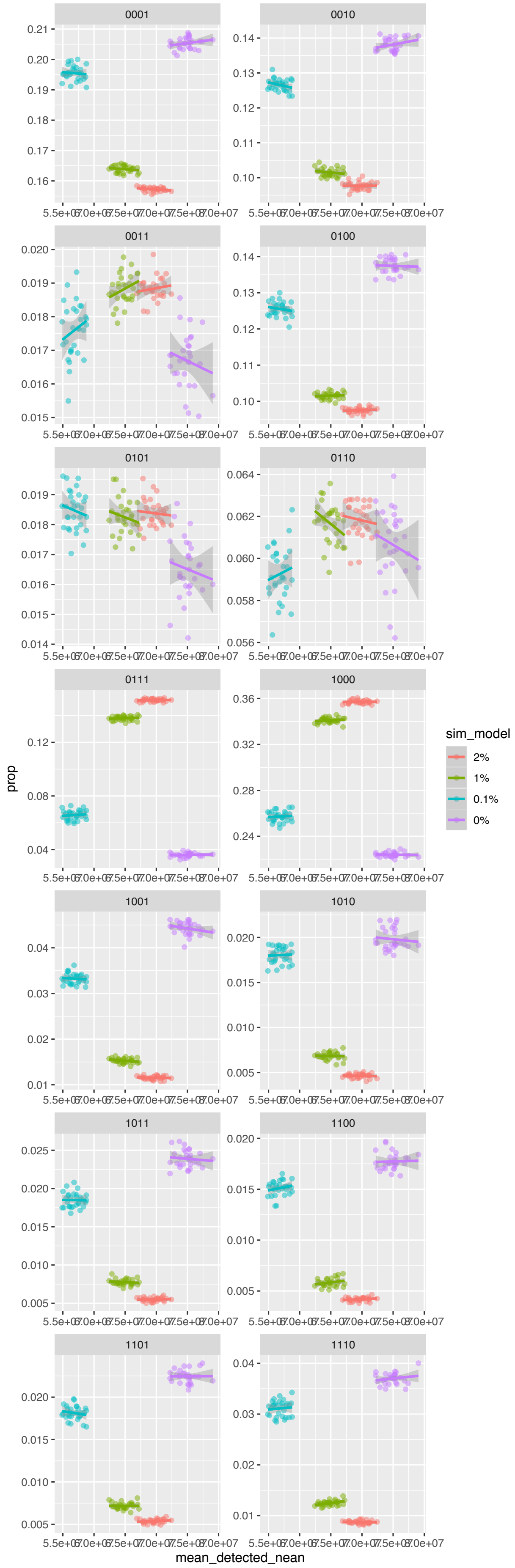


**Fig. S15.** Correlation between detected Neanderthal ancestry and different mutation motifs of the type [X, D, N, H] in simulated data. The proportion of mutation motifs (y-axis) was calculated for each simulated individual in different models of super-archaic introgression. The total amount of Neanderthal is shown in base pairs (x-axis) as detected by *HMM* in the different models. Note that models with no super-archaic introgression always resulted in a higher total detected amount of Neanderthal introgression, in similar amounts to the real data. Once again, we disregarded mutation motifs of the types 0000 and 1111 where all individuals carry the ancestral or derived alleles, respectively.
